# Activity-Dependent Ectopic Spiking in Parvalbumin-Expressing Interneurons of the Neocortex

**DOI:** 10.1101/2024.01.22.576676

**Authors:** Brian B. Theyel, Rachel J. Stevenson, Barry W. Connors

## Abstract

Canonically, action potentials of most mammalian neurons initiate at the axon initial segment and propagate bidirectionally: orthodromically along the distal axon, and retrogradely into the soma and dendrites. Under some circumstances action potentials may initiate ectopically, at sites distal to the axon initial segment, and propagate antidromically along the axon. These ‘ectopic action potentials’ (EAPs) have been observed in experimental models of seizures and chronic pain, and more rarely in nonpathological forebrain neurons. Here we report that a large majority of parvalbumin-expressing (PV+) interneurons in upper layers of mouse neocortex, from both orbitofrontal and primary somatosensory areas, fire EAPs after sufficient activation of their somata. Somatostatin-expressing interneurons also fire EAPs, though less robustly. Ectopic firing in PV+ cells occurs in varying temporal patterns and can persist for several seconds. PV+ cells evoke strong synaptic inhibition in pyramidal neurons and interneurons and play critical roles in cortical function. Our results suggest that ectopic spiking of PV+ interneurons is common, and may contribute to both normal and pathological network functions of the neocortex.

**SIGNIFICANCE STATEMENT:** A form of neuronal firing that emerges in distal axons and terminals – the ‘ectopic action potential’ (EAP) – has been detected in a few cell populations of the cerebral cortex. Previous investigations of parvalbumin-positive interneurons in neocortex had suggested only a small percentage of cells can fire EAPs. We found that a large fraction of parvalbumin-positive interneurons in the superficial layers of neocortex, including first-order and higher-order areas, can fire EAPs. These results broaden our understanding of the intrinsic firing characteristics of these critically important inhibitory interneurons.

## INTRODUCTION

In most healthy vertebrate central neurons, including those of the cerebral cortex, action potentials are widely thought to be triggered at the axon initial segment (AIS) before propagating into the rest of the cell (Bender and Trussell, 2012; Kole and Brette, 2018). In some cases, however, action potentials can originate from ectopic sites, including axon terminals (Pinault, 1995). Once triggered, these “ectopic action potentials” (EAPs) propagate antidromically along the axon and into the soma and dendrites, as well as orthodromically down axonal branches. In vertebrates, EAPs have been observed most commonly in hyperexcitability-related pathological states. For example, they can arise in cortical and thalamic excitatory neurons whose axons terminate in seizure-generating cortical areas (Gutnick and Prince, 1972, 1975; Noebels and Prince, 1978; Pinault and Pumain, 1985; Pinault, 1995; Keros and Hablitz, 2005). Acutely or chronically injured axons can also generate spontaneous EAPs, which may mediate some forms of persistent pain (Amir, 2005; Devor, 2009).

EAPs have also been observed in nonpathological states, in both invertebrate and vertebrate neurons (Pinault, 1995; Bucher and Goaillard, 2011). Studies of crustacean neurons have been especially informative, with data supporting differential functionality of EAPs when compared to standard action potentials originating from the spike initiation zone (Meyrand et al., 1992; Bucher et al., 2003; Daur et al., 2009; Daur et al., 2019). In mammals, spontaneous spikes were detected in peripheral nerves after high frequency nerve stimulation, and confirmed to originate in the motor nerve terminal (Standaert, 1963). EAPs have also been observed in certain neurons of mammalian cerebral cortex. Distal axons of excitatory pyramidal neurons in hippocampal areas CA1 and CA3, for instance, initiate action potentials linked to sharp wave-ripple complexes and gamma rhythms; these may be triggered in distal axons by a GABA_A_ receptor-mediated mechanism (Papatheodoropoulos, 2008; Bahner et al., 2011; Dugladze et al., 2012). Pyramidal neurons of the mouse neocortex can generate relatively sparse EAPs following stimulation of their somata (Zhang et al., 2023).

There have been several reports of EAPs in inhibitory interneurons of cerebral cortex. In a notable series of studies, neuropeptide Y (NPY)-expressing interneurons of the hippocampus robustly and reliably generated prolonged “barrages” of EAPs in response to somatic depolarization (Sheffield et al., 2011; Sheffield et al., 2013). These barrages lasted for up to tens of seconds and were thought to be mediated, at least in part, by biochemical interactions between the presynaptic terminals of axons and nearby astrocytes (Deemyad et al., 2018). NPY neurons comprise a small proportion of inhibitory cells in the hippocampus, but the robustness of ectopic spikes detected in these studies suggests they may play an important role in hippocampal function.

The most common type of inhibitory cell in cerebral cortex is the fast-spiking, parvalbumin-expressing interneuron (PV+ cell) (Rudy et al., 2011). Studies of EAPs in cortical PV+ interneurons are few and mixed. While many PV+ cells of the hippocampus (Elgueta et al., 2015) and piriform cortex (Suzuki et al., 2014) could be induced to generate EAPs, a study of primary visual neocortex reported only a single example (Imbrosci et al., 2015), and another study in mouse prefrontal cortex identified EAPs in 10/22 (45%) of cells (Elgueta et al., 2015).

While investigating the intrinsic properties of PV+ cells in mouse orbitofrontal neocortex (OFC), we noticed frequent ectopic spiking, which led us to further explore its prevalence and properties in both OFC and the barrel field of primary somatosensory cortex (S1BF). Here we demonstrate that the majority of the PV+ interneurons in supragranular layers of the mouse OFC and S1BF can fire trains of EAPs and that nearly all can generate at least one EAP. Thus, EAPs may play a much more prominent role in neocortical processing than previously appreciated.

## RESULTS

We made somatic whole-cell recordings from neocortical PV+ and somatostatin-expressing (SOM+) interneurons in acute sagittal slices of mouse OFC (Fig. 1), or from S1BF cut in the thalamocortical plane (Agmon and Connors, 1991). Most of the data reported here were obtained from PV-Cre x TdTomato or SOM-Cre x TdTomato transgenic mouse lines. Results were obtained from 35 male and female mice between the ages of P21 and P60. Fig. 1A,B illustrates fluorescent images of PV-TdTomato expression from OFC slices, and highlights the ventral orbital (VO) and lateral orbital (LO) regions where recordings were made from neurons in layer 2/3 (unless otherwise indicated). Fig. 1C shows a representative PV+ interneuron of layer 2/3 of VO; its axon densely ramifies both locally and laterally within layers 1 and 2/3. Note that all data reported here were from neurons in layer 2/3.

**Figure 1:**
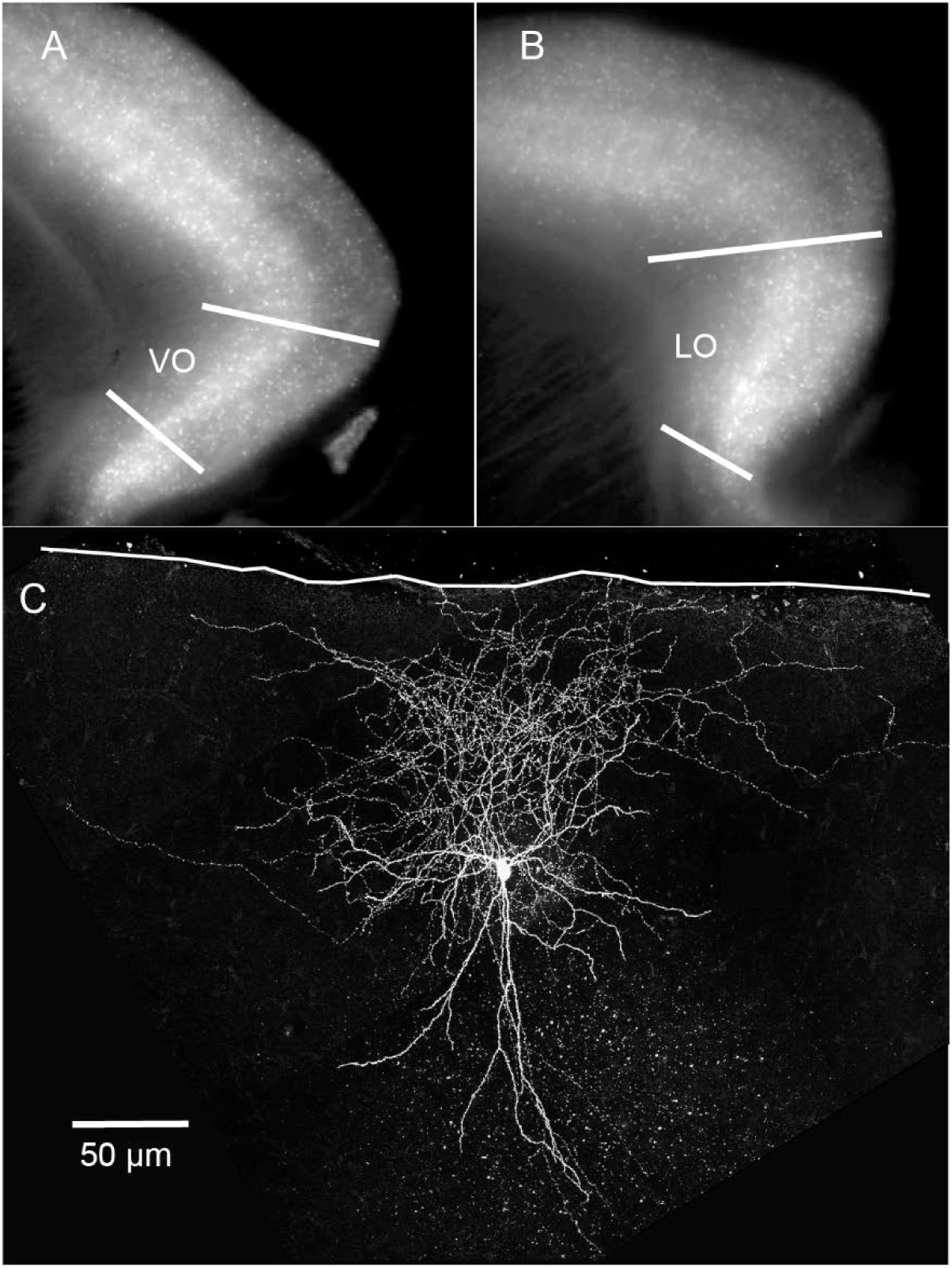
Anatomical location of OFC recordings and filled PV+ cell. A) and B) Mouse brain slices from a PVCre x Ai14 mouse under fluorescence illumination C) Confocal stack of a PV+ cell recorded in area VO. Cell filled with biocytin during recording and immunolabeled with a fluorescent dye. White line = pia. VO = Ventral orbital cortex; LO = Lateral orbital cortex.

### Basic phenomenology of ectopic action potentials in PV+ interneurons

We first noticed atypical action potentials in somatic recordings from layer 2/3 PV+ cells of OFC while running a protocol designed to measure action potential frequency as a function of stimulus current intensity. We found that initial sweeps of sequential injections of current steps triggered the expected trains of conventional action potentials during the stimulus pulses (Fig. 2A, top two traces; current steps were 600 ms long, increasing by 5 pA increments, applied every 2 sec until each cell either fired a sustained train of EAPs or reached depolarization block). At a certain point in the protocol, usually after hundreds to just over one thousand action potentials had been triggered (see below and Supp. Table 1 for quantification), most recorded neurons began to fire action potentials persistently after current steps ended (e.g. Fig. 2A, lower traces). These action potentials differed from evoked action potentials in that they rapidly rose from resting membrane potential, well below spike threshold, and their amplitudes often varied markedly. The waveform of these action potentials strongly suggests that they propagated antidromically, appearing in the somatic recording without the gradual depolarizing phase representing charging of the somatodendritic compartment by summating post-synaptic potentials (Pinault, 1995).

**Figure 2:**
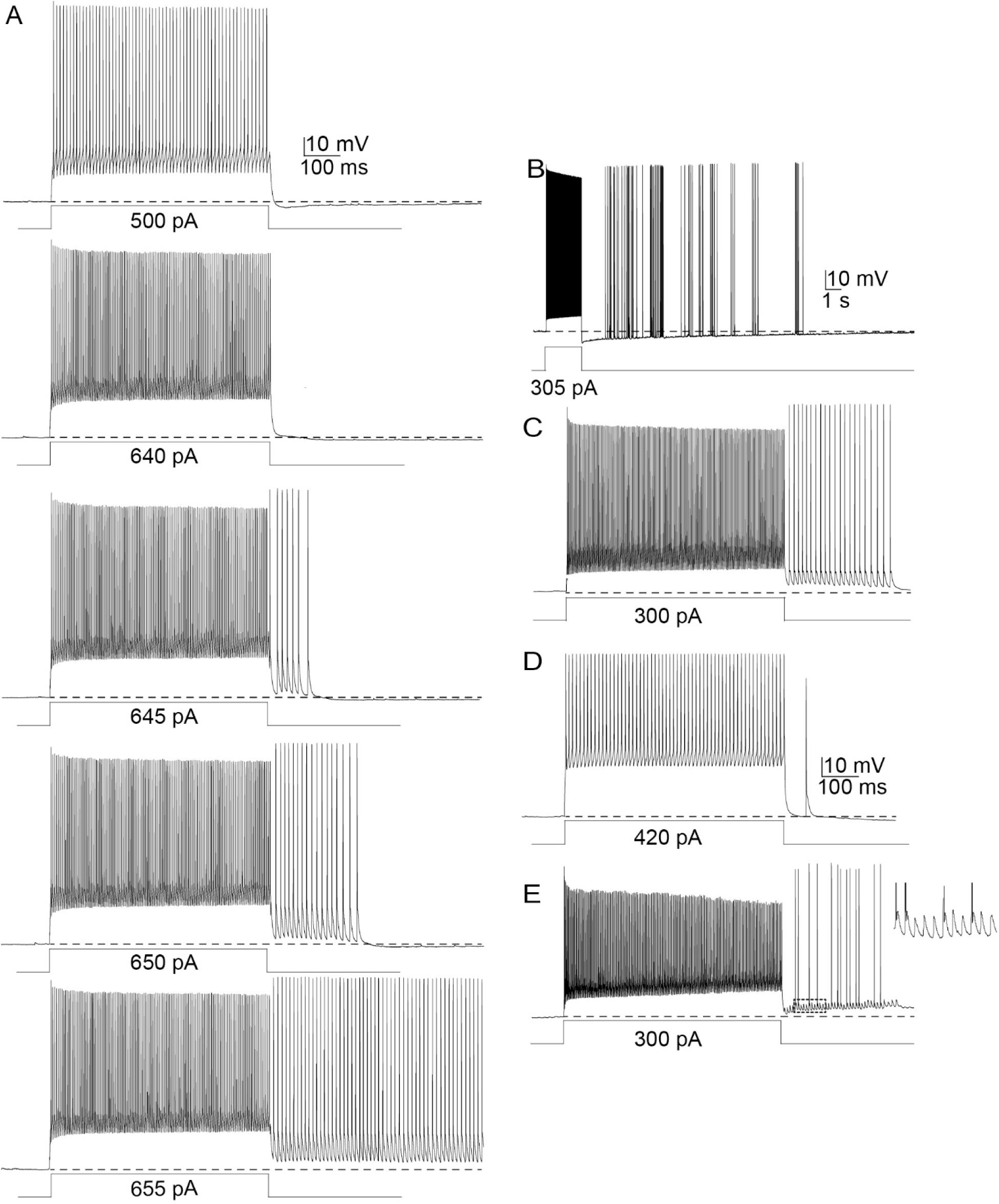
Different patterns of ectopic action potentials in PV+ cells. A) Example of induction of EAPs in a single PV+ cell. The protocol consisted of 600 ms long positive current steps, increasing in amplitude by 5 pA every 2 second sweep. B-D include data from 4 separate cells. B) Sporadic, scattered ectopic spiking lasting several seconds. C) Large amplitude EAPs. D) Single medium-amplitude EAP. E) Mixture of high and small-amplitude EAPs. Right-most trace is a magnification of the area outlined in a dotted line in the left trace. All recordings are in the whole-cell current clamp configuration. Note that all action potentials after the current steps are EAPs.

These post-stimulus antidromic action potentials are reminiscent of “backfiring” action potentials, recorded *in vivo* and *in vitro* in multiple species. Action potentials with these properties, which initiate in the distal axon, have often been observed in mammalian models of epilepsy (Gutnick and Prince, 1972; Gutnick and Prince, 1974; Noebels and Prince, 1978; Pinault and Pumain, 1985; Stasheff et al., 1993; Pinault, 1995), and in normal crustacean neurons (Meyrand et al., 1992; Bucher et al., 2003; Bucher and Goaillard, 2011; Daur et al., 2019). They also resemble the action potentials noted and described more recently as “barrage firing”: spontaneous, ectopically initiated, back-propagating action potentials recorded from somata of several types of inhibitory interneurons in cerebral cortex, usually following extended somatic stimulation (Sheffield et al., 2011; Suzuki et al., 2014; Elgueta et al., 2015; Imbrosci et al., 2015; Rózsa et al., 2023). Here we have elected to continue to use the term ‘ectopic action potentials’ to refer to action potentials that appear from their physiologic signature to originate in distal axonal sites.

A variety of stimulus protocols elicited EAPs in PV+ cells of the OFC. These included repeated trains of brief (0.1 msec) suprathreshold pulses of injected current at varying frequencies and durations, optogenetic (ChR2) activation, and prolonged (0.5-1 sec) suprathreshold current pulses repeated at 0.3-3 Hz with steady or incrementally increasing amplitudes. The latter protocol, with incrementing (by 5 pA) current steps lasting 600 ms every 2-3 seconds, was the most effective and practical; it is also similar to protocols used previously on other neuron types (Sheffield et al., 2011; Sheffield et al., 2013; Deemyad et al., 2018), and we used it to obtain the data described below unless stated otherwise.

We were able to elicit ectopic spiking in nearly all sampled PV+ neurons of OFC (43/46 cells; 93%). Ectopic spiking typically began after at least several hundred orthodromic action potentials were triggered by somatic current injection. It took 1,236 ± 772 (mean ± standard deviation throughout unless otherwise noted) triggered action potentials (median 989) over 70.1 ± 46.4 seconds to elicit the first EAP in OFC PV+ cells that fired EAPs. Ectopic spiking was not limited to the OFC; 100% (21/21) of sampled PV+ cells in layers 2/3 of S1BF fired EAPs. In S1BF, it took an average of 1,516 ± 1679 action potentials (median 997) to elicit the first EAP. Combining data from the two areas (OFC and S1BF), 64/67 PV+ cells (96%) fired EAPs.

To decrease the probability that the EAPs we detected were generated by network activity, we applied DNQX and APV to block fast excitatory neurotransmission in a subset of experiments. We found that 11/12 (91.7%) recorded neurons fired EAPs in the presence of AMPA and NMDA receptor blockade, suggesting that EAP initiation is independent of fast glutamatergic transmission and surrounding network activity.

The patterns of ectopic spiking in PV+ cells varied from spike barrages that persisted for several seconds (Fig. 2A,B-E) to as few as one EAP. We define a “barrage” as a train of EAPs beginning within 50 ms of the end of stimulus current pulses and lasting at least 250 ms, with a mean frequency of at least 50 Hz, similar to criteria used by others (Sheffield et al., 2011). Ectopic spike trains that did not meet criteria for a barrage were classified as ‘other.’ We have distinguished between barrage firing and other EAP firing here to better compare our findings to studies of other cell types that quantified the prevalence of “barrage” (or “persistent”) firing and EAPs that did not meet barrage criteria (e.g. Sheffield et al. 2011). Fig. 3 depicts spike activity from all sampled neurons as a raster plot. One trial from each neuron was chosen to represent its maximal EAP activity: the stimulatory sweep during which that neuron fired the most EAPs. Among the OFC PV+ cells tested (Fig. 3A), 74% (34/46) generated full barrages of EAPs, 20% (9/46) fired multiple EAPs but did not reach our criteria for a barrage, and 2% (1/46) fired only a single EAP; 7% of the cells (3/46) fired no EAPs. In S1BF, 13/21 (62%) recorded PV+ cells fired full barrages of EAPs, and 8/21 (38%) PV+ cells fired multiple EAPs but fell short of criteria for a full barrage (Fig. 3B).

**Figure 3:**
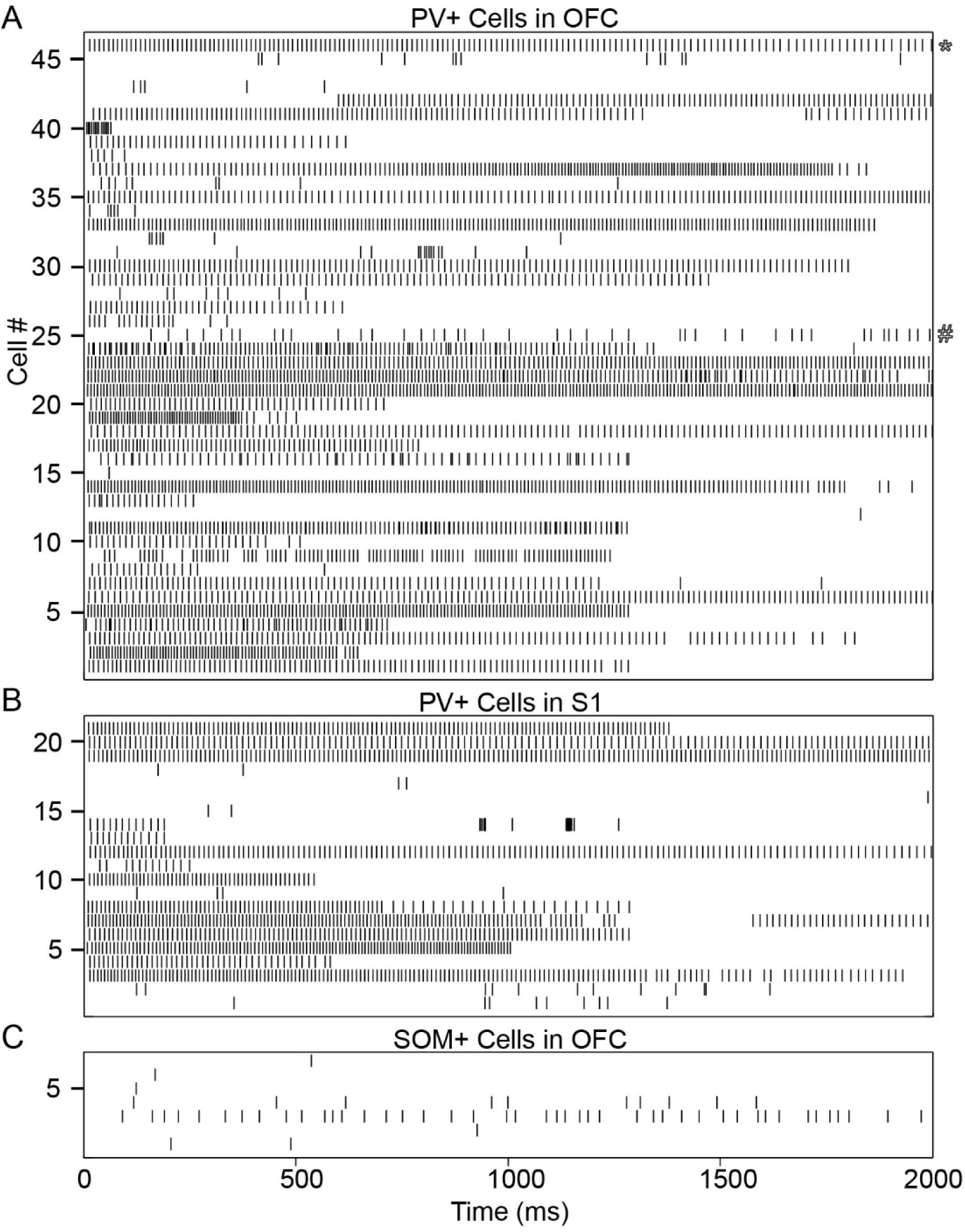
Spike raster of the first two seconds of ectopic firing in all recorded cells that fired ectopic action potentials. A) Ectopic spike raster plot of PV+ cells in OFC. B) PV+ cells in S1BF. C) SOM cells in OFC. Action potentials were detected using PClamp, with manual verification of accuracy. The starred raster (top) is an example of a train of EAPs exhibiting spike frequency adaptation, and the pound sign is next to an example of a cell with a wide range of interspike intervals.

Patterns of ectopic spike trains varied from cell to cell. Spike trains, as shown in the spike time histogram in Fig. 3, often exhibited spike frequency adaptation, and often consisted of clusters of spikes separated by gaps. The frequency of ectopic firing in each cell was calculated using the interspike interval (ISI) between the second and third ectopic action potential. The mean frequency of ectopic firing across PV+ cells was 91.85 +/- 24.89 Hz. Spike frequency adaptation was measured in the 34 cells with at least 10 EAPs by taking the mean of the last five ISIs and dividing it by the mean of the first five ISIs to create an adaptation ratio. The mean and SD of this adaptation ratio was 1.48 +/- 0.78, with a range from 0.75 to 4.67, suggesting that trains of ectopic spikes decrease in frequency over time, but with some exceptions. Variability in spike frequency was measured using a version of the coefficient of variation (CV_2_) that is not as influenced by the large ISIs sometimes found in our dataset as other measures of spike variability are (Holt et al., 1996). CV_2_ ranged from 0.02 to 0.39, with a mean and SD of 0.12 +/- 0.09 across cells, indicating relatively low variability in spike frequency during ectopic firing.

### Ectopic spiking in chandelier cells and somatostatin positive cells

PV+ / fast-spiking interneurons can be divided into multiple subtypes. The best characterized subtypes of PV+ cells are basket cells and chandelier cells (also known as axo-axonic cells), which are morphologically distinct and synapse upon different postsynaptic compartments (Miyamae et al., 2017). Basket cells target the soma and proximal cell body of pyramidal cells and other interneurons; chandelier cells target the proximal axon of pyramidal cells with abundant synaptic boutons arranged vertically (called ‘cartridges’) (DeFelipe et al., 2013). This differential targeting yields differences in function: postsynaptic responses of chandelier cells can be either excitatory or inhibitory depending on network state (Szabadics et al., 2006; Woodruff et al., 2009; Woodruff et al., 2011), whereas basket cell output is more generally inhibitory. Thus, EAPs, which are likely to occur during periods of network excitation, might have different roles in these cell types.

We recorded from six physiologically identified chandelier cells in OFC, all of which fired EAPs. Five of them fired full amplitude EAPs, and one cell fired small amplitude EAPs. Five cells fired sufficient EAPs to meet our criteria for a barrage, and one cell fired a single EAP. It took an average of 1,680 +/- 1,253 evoked action potentials to generate the first EAP in these cells. While our sample is too small for meaningful statistical comparison, chandelier cells are qualitatively similar to basket cells in both the number of spikes required to induce EAPs and in EAP firing patterns once elicited.

In an exploratory analysis we attempted to elicit EAPs in SOM+ cells. We found that 88% (7/8) of recorded SOM interneurons in the OFC fired EAPs. No SOM+ cells met our criteria for full barrages; 5/8 cells fired multiple ectopic spikes, and 2/8 fired single ectopic spikes on their most active runs (Fig. 3C). See supp. Table 1 for further details.

### The shapes and sizes of ectopic action potentials

As action potentials propagate via excitable membranes of varying geometry and excitability—e.g. along axons, through the AIS, into the soma, through the dendritic arbor—their shapes and sizes usually change in significant and informative ways (Eccles, 1957; Ramón et al., 1976; Debanne et al., 2011). Here we describe in more detail the spike morphologies of the EAPs we recorded from somata of PV+ cells.

The amplitudes of the smallest EAPs detected during our somatic recordings ranged from about 2-6 mV, defined here as “small-amplitude” (e.g. Fig. 2E). Less commonly, some neurons generated medium-amplitude EAPs that ranged from 10 mV to 80 mV in height (e.g. Fig. 2D). In some PV+ cells, EAPs reached peak voltages more positive than those of the current-triggered action potentials that preceded them (Fig. 2A,B; Fig. 4A,B). The peaks of the largest EAPs ranged from about 90-110 mV above resting membrane potential, which was comparable to or larger than the first action potentials of current-triggered spike trains; 25/69 (36.2%) of PV+ cells in OFC and S1 fired EAPs that were larger than their evoked counterparts. For comparison, non-ectopic spike heights averaged 93.0 +/- 12.9 mV. In some PV+ cells EAP amplitudes were distinctly bimodal, fluctuating between large- and small-sized over the course of the recording (see Fig. 2E). For descriptive purposes, large-sized EAPs were defined as those at least 90% of the amplitude of the first spike in the evoked spike train. Medium-amplitude EAPs were defined as those between 10% and 90% of the first evoked action potential, and small-amplitude EAPs were <10% of evoked spike height. Fig. 5 shows the pooled distribution of EAP amplitudes recorded from PV+ cells in OFC during sweeps in which the most EAPs occurred in any given cell. Among our sample of 43 PV+ cells that fired EAPs in OFC, 24 cells fired only large EAPs, 1 fired only medium EAPs, 2 fired only small-height EAPs, 10 fired both small-amplitude and large-amplitude EAPs, 5 cells fired both medium and large EAPs, and 1 cell fired medium and small EAPs. SOM+ neurons generated only large-amplitude EAPs.

**Figure 4:**
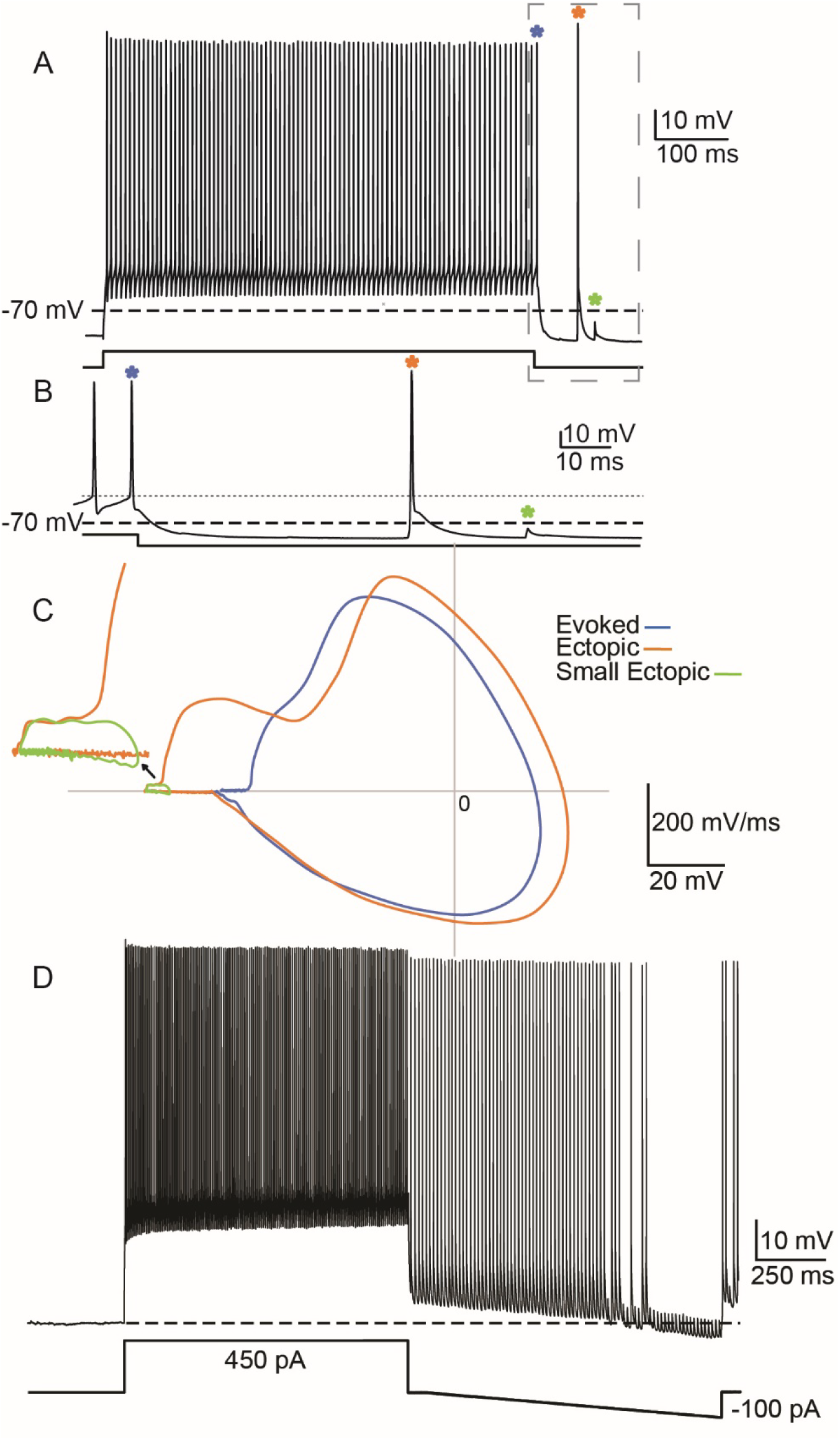
Evoked and ectopic action potentials are qualitatively different: A) Zoomed out trace of evoked action potentials. B) Zoomed in portion of the trace in panel A. Note the classic rise to threshold in the evoked action potentials on the left, and the immediate steep rise without a readily apparent lower-slope rise to threshold in the EAP to the right. C) Corresponding phase plots (dV/dt vs. membrane potential). The orange line corresponds to the large amplitude EAP shown in A and B (marked by orange asterisk) and the green and blue lines are the small amplitude (small) EAP and last evoked action potential, respectively. The green inset depicts a ‘zoomed in’ version of the short ectopic to better illustrate its shape. D) Large-amplitude ectopic spiking can be reduced to small-amplitude with hyperpolarizing current. Here, I stimulated a PV+ cell by injecting 1 second 450 pA current steps followed by 100 ms with no current injected, followed by a 1-second hyperpolarizing current ramp that gradually decreased from 0 down to -100 pA. The above trace shows a run of EAPs that transition from large-amplitude to small-amplitude action potentials, presumably due to failure of propagation beyond a node of Ranvier or an axonal branch point, when I hyperpolarized the cell during a run of EAPs.

**Figure 5:**
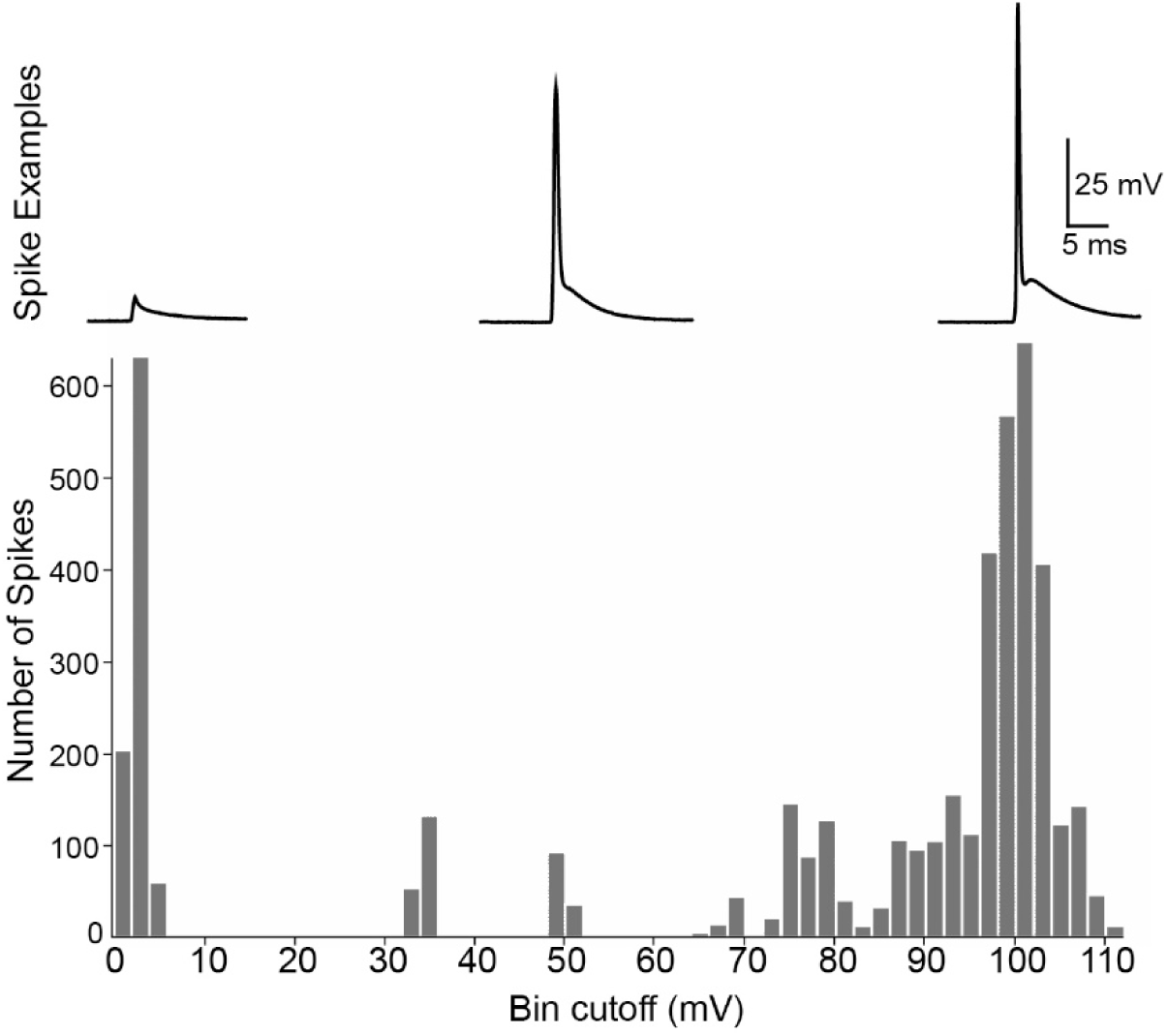
Distribution of ectopic spike amplitudes. EAP distribution in PV+ cells in OFC. This is a pooled distribution of all EAPs that we elicited from every recorded PV+ cell. Bin width was 2 mV. Note bimodality at small and high-amplitudes with wide variability of spike heights greater than 10 mV and less than 80 mV.

One possible explanation for the lower amplitudes of small- and medium-amplitude EAPs is that they represent axonal spikes that fail to fully propagate to the cell body, and are interrupted somewhere between the site of initiation and the recording electrode located at the soma (Coombs et al., 1957; Moore et al., 1983; Sheffield et al., 2011). Full or partial failure of propagation can occur at sites of abrupt impedance mismatch (e.g. axonal branch points, axon-soma transitions), reduced excitability (e.g. due to region-specific ion channel expression), or local membrane hyperpolarization. We tested the latter condition by inducing EAPs in the usual way but followed the depolarizing current pulse with a hyperpolarizing current ramp to a minimum of -100 pA (similar to Sheffield et al., 2011) in two cells. During the hyperpolarization, large-sized ectopic spikes switched to small-amplitude ectopic spikes (Fig. 4D); when the negative ramp current terminated, the action potentials reverted to a mix of large- and small-amplitude.

The shapes and dynamics of somatically recorded action potentials are also informative. Ectopic and evoked action potentials have qualitatively different shapes, which is particularly evident in their phase plots (see Fig. 4A-C). The most striking difference is reflective of the different thresholds of ectopic spikes: the value of dV/dt is high at membrane potentials below the cells’ evoked action potential thresholds. When the membrane potential reaches threshold during an ectopic action potential, there is a further increase in slope before reaching the peak dV/dt at approximately the same point as evoked action potentials. Small amplitude EAPs also had a characteristic phase plot and did not reach threshold for evoked action potentials (see green asterisks and trace in Figs 4A-C).

### Post-stimulus shifts of somatic membrane potential and ectopic spike generation

Some PV+ cells remained slightly depolarized for a brief period after our current injections had ended (∼0.5-5 mV above resting potential; e.g. Fig. 6B,C). These depolarizations immediately followed the stimulus current steps and were often terminated abruptly by afterhyperpolarizations (AHPs; Fig. 6B). Occasionally one or more waves of subthreshold depolarization followed the initial depolarization and its AHP (Fig. 6C). More examples of prolonged subthreshold depolarizations are shown in Figs. 2A (bottom 4 traces), 2C, 2E, 6B-C and 7E-H. While not conclusive, the coincident timing of these depolarizations and ectopic spike generation suggests a causal relationship: small-amplitude depolarizations recorded at the soma may be passively propagated large depolarizations that initiate ectopic firing at distal axonal sites. In effect, the recording electrode at the soma may detect depolarizing waves underlying EAPs when they’re generated nearby, but only the (actively propagated) EAPs are measurable when they are initiated far from the soma. The fact that we observed EAPs riding on these depolarizations sometimes (Fig. 6B,C), but not in all cases (Fig. 6A), may reflect wide variability in the electrotonic distances between the soma and the sites of ectopic spike initiation. Slow activity-dependent potassium conductances and the AHPs they mediate could also obscure slow depolarizations at distal axonal regions.

**Figure 6:**
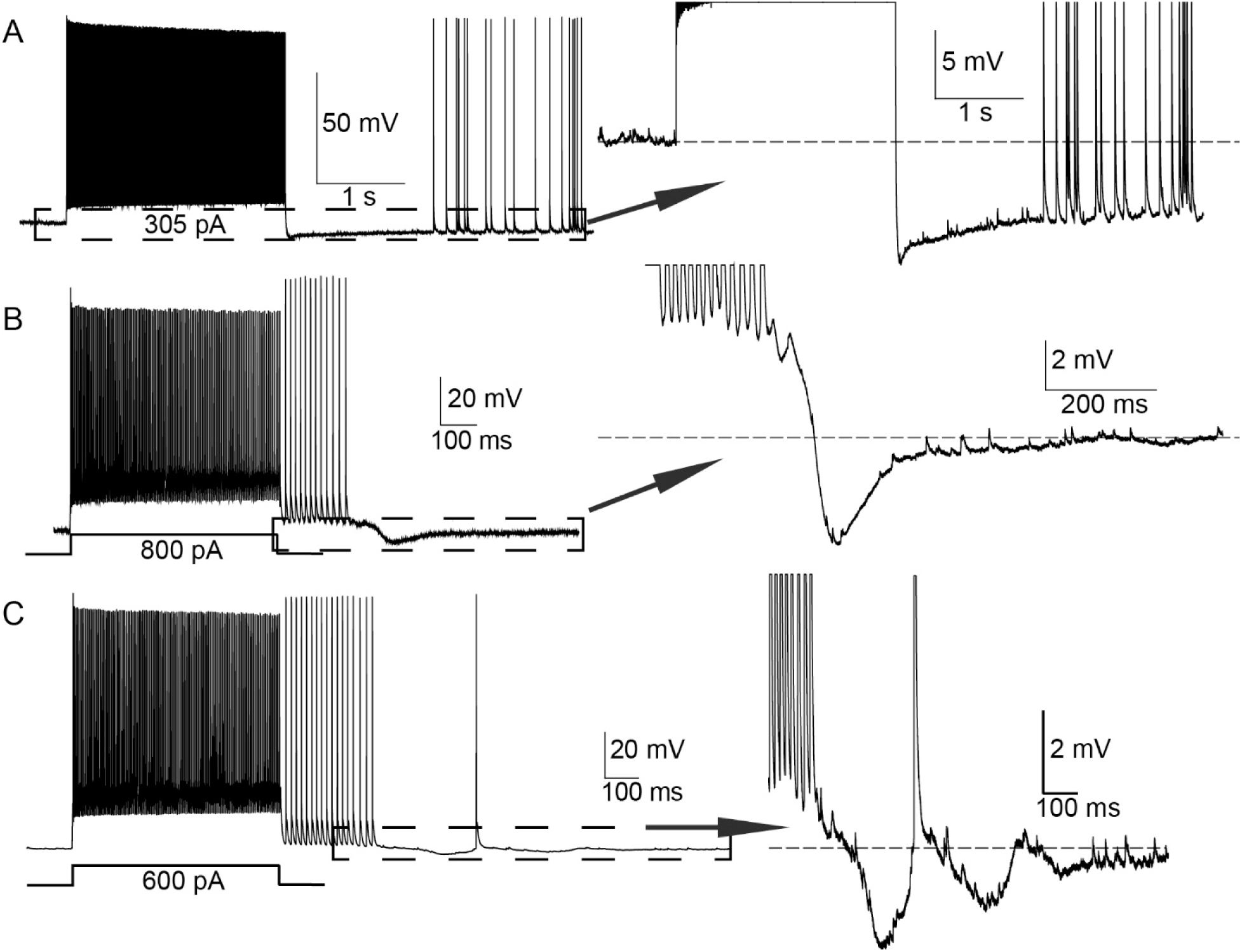
Ectopic spikes emerge from variable baseline membrane potentials. Traces from three different cells are shown in A-C. A) PV+ cell showing an example of ectopic spikes emerging during hyperpolarization (relative to resting membrane potential). B) Ectopic spike train emerging during a wave of depolarization that immediately follows the evoked (with the shown current step) spike train. C) Cell in which ectopic firing emerged during waves of depolarization with an oscillatory component. Dotted line boxes depict the portion of the traces shown at magnified scales in the inserts.

### Multiple ectopic action potential initiation sites in single cells?

EAPs are generated in the axon, its branches, and/or its presynaptic terminals. It is possible that ectopic spikes can be initiated at multiple locations in the axonal arbor of an individual neuron. There are at least four ways this might be detected in whole-cell somatic recordings: First, when EAPs initiate electrotonically close to the cell body, waves of depolarization that trigger ectopic spikes might be recorded. If ectopic spike generation is electrotonically distant, however, these depolarizations might be too small to detect with somatic recordings. Thus, if some EAPs are observed to be ‘riding’ on waves of depolarization while other EAPS in the same recording are not, this may suggest two sources of ectopic spiking in a single neuron. Second, axonal branch points may serve as areas where backpropagating spikes fail due to impedance mismatch, manifesting as ‘medium’ or ‘small’ amplitude action potentials as recorded at the soma. In this case, detecting spikes with distinct heights in the same cell could suggest two sources of ectopic spike generation. Third, one might imagine a different threshold for ectopic spike initiation at two different sites, which would manifest as ectopic spiking arising after different numbers of action potentials occur over a certain amount of time within the same cell. Fourth, EAPs might emerge from two different sources simultaneously, which would manifest as co-occurring ectopic spiking at different frequencies. We will highlight evidence from two cells supporting the idea that EAPs can emerge from two or more distinct distal sites.

*Cell 1:* Some evidence for EAPs arising from different sites within an individual cell is presented in Fig. 7A1-8. Early in the stimulus protocol (Fig. 7A2-4) the cell generated large-amplitude ectopic spikes that emerged from a baseline near the cell’s resting membrane potential. These EAPs emerge 1,675 action potentials (elicited over the course of 36 seconds) before the sequences of short-amplitude EAPs depicted in Fig. 7A5-8. Note that these short-amplitude ectopic spikes were only present during prolonged, small depolarizations that were either fairly steady (Fig. 7A5-8) or oscillatory (Fig. 7A7). In this cell, two sets of EAPs were qualitatively different in three ways: they had different activation thresholds (the number of evoked spikes required to trigger them), they exhibited different amplitudes, and one set arose from a hyperpolarized baseline while the other set occurred during waves of depolarization.

**Figure 7:**
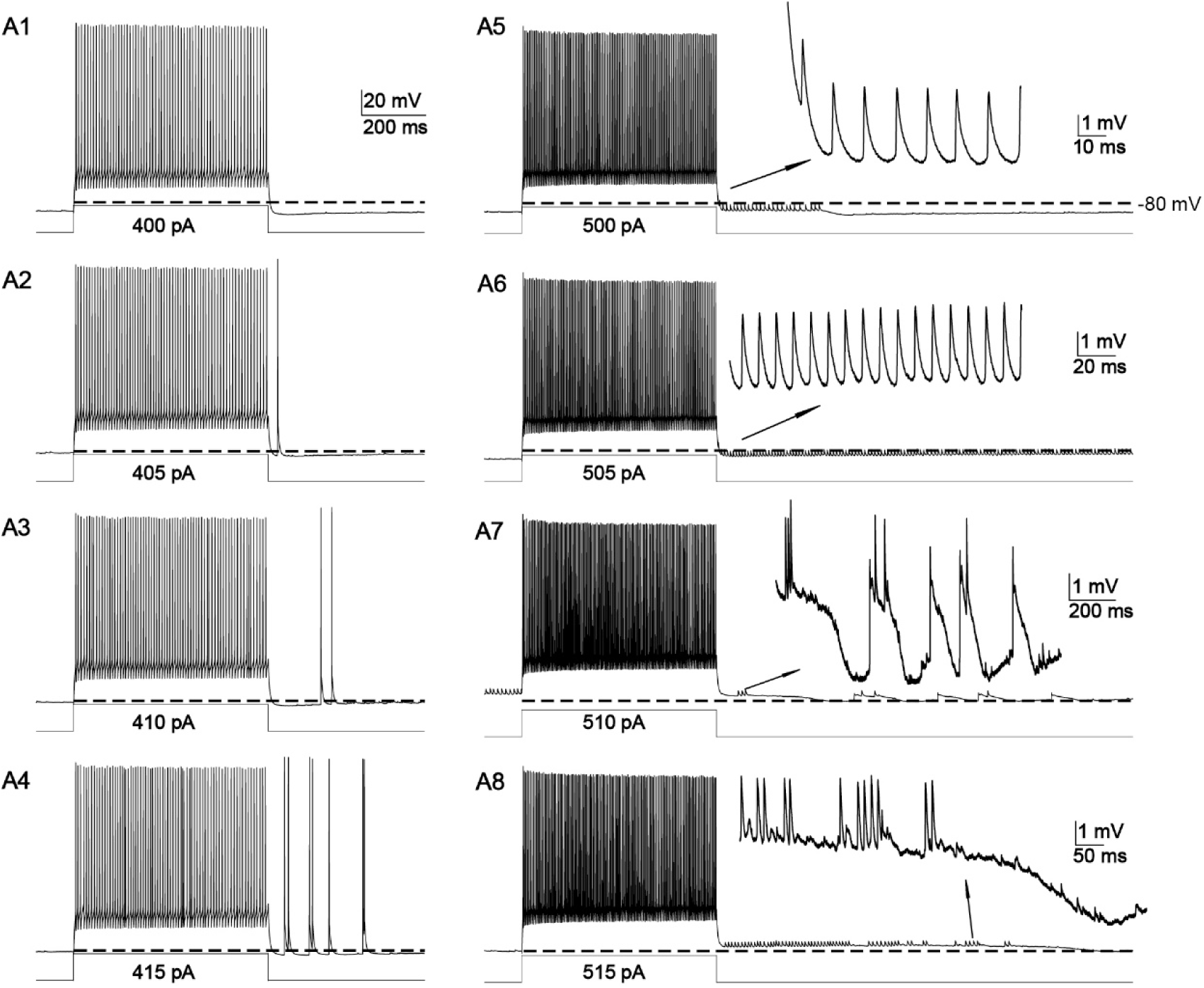
Ectopic action potentials from two sources: occurrence within and outside of a long, low-amplitude period of depolarization. Traces from one cell are depicted in A1 to A8; Traces from another cell are shown in B1 to B2. A1-A4) High amplitude EAPs evoked over subsequent runs; note that they emerge from membrane potentials that are slightly hyperpolarized relative to baseline (horizontal dotted lines). A5-A8) Small EAPs emerging when membrane potentials are above baseline. Scale bar in A1 applies to zoomed out traces in A1-A8. Inset scale bars in A5-A8 correspond to zoomed in depictions of small-amplitude EAPs.

*Cell 2:* In this cell (Fig. 8), small- and large-amplitude EAPs co-occurred before a period of time during which small EAPs persisted. While not all of the small EAPs were visualized during the period that both small and large EAPs were visualized (first ∼250 ms), when they were both present small EAPs preceded large EAPs by a significantly shorter time period than they followed large EAPs. The average time between small EAPs and subsequent large EAPs was 2.3 +/- 1.1 ms, whereas the time between small EAPs and the preceding large EAPs was 10.6 +/- 1.1 ms (2-tailed t-test p-value 5.4 x 10^-9^). A 2.3 ms interspike interval is equivalent to a firing rate of 435 Hz. Given that this is close to the maximal PV+ firing rate of ∼450 Hz it is unlikely that both the large and small EAPs originated from the same site. This suggests that the small EAPs were triggering large EAPs. Further evidence for two sources of ectopic firing is that when both small and large EAPs were co-occurring they appeared to ride on a wave of low amplitude depolarization that partially repolarized after the large EAPs ceased and small EAPs persisted. If we assume that this wave is a readout of depolarization in an electrotonically nearby site, where the large ectopic spikes are being generated, the persistence of the short ectopic spikes after partial repolarization again suggests an origin at a separate, electrotonically distant site. A more distant site is also more likely to have points at which EAPs would fail to backpropagate (e.g. at a branch point), which may help to explain the spike amplitude difference. In this cell, two sets of ectopic spikes were qualitatively different in two ways: they were of different height, and one set rode on a wave of depolarization that partially repolarized while the other set persisted (see Fig 8). This suggests that one site generating ectopic spikes stopped doing so while the other persisted.

**Figure 8:**
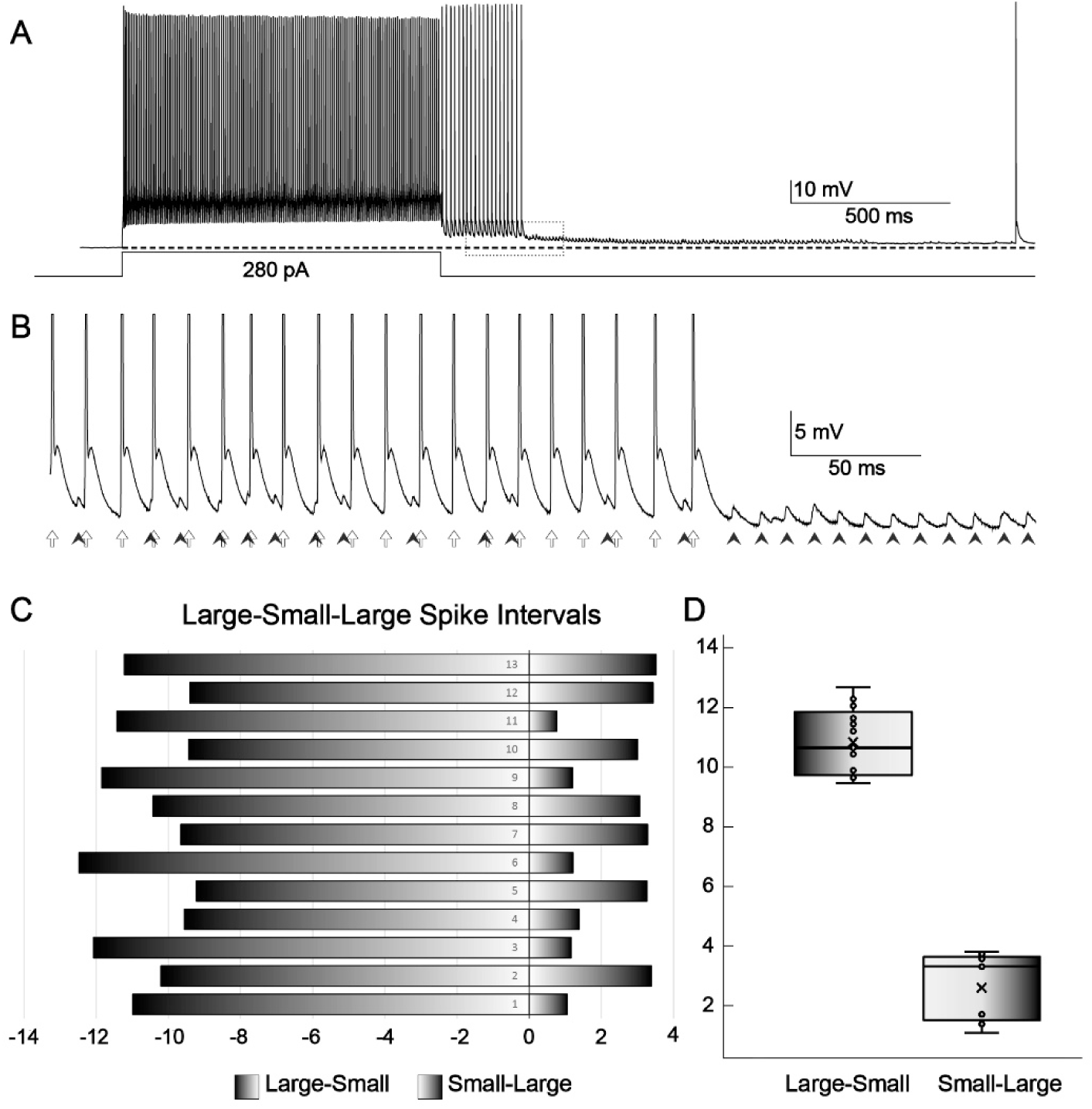
Co-occurrence of large and small EAPs suggesting two sources. **A**) Trace from a cell with both high- and small-amplitude EAPs. B) Zoomed in area bounded by dotted line box in A. Note that small-amplitude EAPs interspersed between high-amplitude EAPs are at a different frequency from large-amplitude EAPs, suggesting that they arise from different sources within the axonal arbor. C) Interspike intervals from large spikes to small spikes (negative numbers) and small spikes to subsequent large spikes (positive numbers), in ms. D) Plot of intervals between large and small spikes. The intervals from preceding large spike to small spikes are on the left, and from small spikes to following large spikes are plotted on the right.

### Synaptic output triggered by ectopic action potentials

We observed the synaptic effects of ectopic spikes during paired recordings from a PV+ cell and a regular-spiking (RS; presumed excitatory) neuron in layer 2/3 of OFC. Inhibitory postsynaptic potentials (IPSPs) were recorded in current clamp while injecting sufficient steady current (250 pA) to depolarize the RS cell membrane to approximately -56 mV, well under the cell’s spike threshold of -48 mV. Fig. 9 illustrates two ectopic spikes from the PV+ cell (top) and two short-latency IPSPs recorded from the RS cell (bottom). The first four postsynaptic potentials (PSPs) evoked during EAPs yielded IPSPs of -0.5 to -0.9 mV. These results establish that ectopic firing can lead to postsynaptic responses; further work will be required to quantify the functional impact of ectopic spike-generated inhibition.

**Figure 9:**
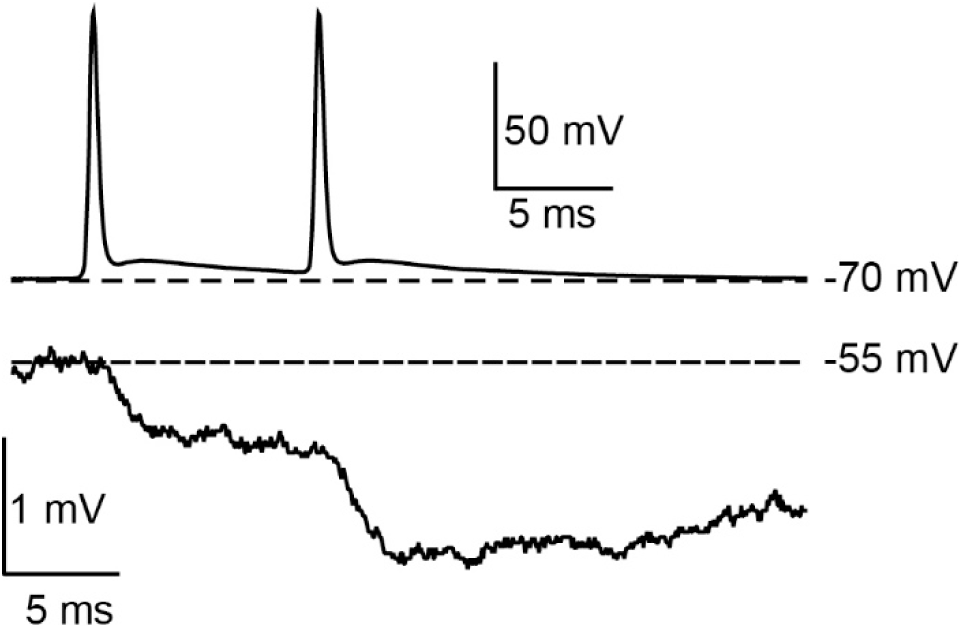
Synaptic output during ectopic action potentials. The top trace is from a PV+ cell during ectopic firing. The bottom trace is a simultaneous paired cell current clamp recording from a regular spiking cell. Dotted lines represent membrane potential in both cells immediately prior to the EAPs emerged. Note that the bottom cell was held near -55 mV with 250 pA of current to bring the cell far enough from chloride’s reversal potential to detect inhibitory inputs.

### Effect of age on ectopic spiking

To determine whether neurodevelopmental changes in PV+ cells alter EAP generation, we analyzed the correlation between age and the threshold for EAP generation. Age had a statistically significant effect on ectopic spike threshold (defined as the number of action potentials necessary to elicit ectopic spiking) of PV+ cells in OFC, as demonstrated by regression analysis (Pearson r = -0.29; P = 0.018). EAPs were more readily elicited in older animals.

## DISCUSSION

The most surprising result of our study is that EAPs—spikes initiated distal to the axon initial segment—could be readily generated in a large majority (>90%) of sampled PV+ interneurons in two different areas of the neocortex. This is a significantly higher proportion than previously reported. Neocortex has been only sparsely studied; Imbrosci et al. (2015) induced ectopic spikes in only one of 24 (∼4%) fast-spiking (presumed PV+) cells in visual neocortex. Elgueta et al. (2015) found that 10/22 (∼45%) of PV+ cells in prefrontal cortex fired EAPs, which is approximately one half of the proportion we observed in both OFC and S1BF.

Prior to this report, it was thought that only a subset of archicortical PV+ cells were highly likely to fire EAPs. Suzuki et al. (2014) reported barrage firing in 23% of fast-spiking, GAD67-GFP-expressing interneurons (presumed PV+ cells) of piriform cortex. Krook-Magnuson et al. (2011) found ectopic spikes in <20% of PV+ cells in the CA1 area of hippocampus. Elgueta et al., 2015 reported that unique PV+ cells—soma- and axon initial segment-targeting interneurons—in the hippocampal dentate gyrus reliably generated EAPs (58-85% of tested cells, depending on the stimulus protocol).

It is not clear why the estimated prevalence of EAPs in PV+ cells ranges so widely across studies. There are many possibilities. The sampled neurons have been drawn from disparate regions of neocortex, archicortex, and paleocortex. Criteria for defining PV+ cells have been inconsistent across studies. Even within individual areas of cortex there are multiple subtypes of PV+-expressing neurons (Tremblay et al., 2016; Pelkey et al., 2017; Gouwens et al., 2020). No study of EAPs has systematically compared PV+ cell subtypes across cortical areas using the same approach. The mouse strains, ages, tissue preparation methods, recording conditions, and stimulus protocols have differed. The propensity for neurons to generate EAPs depends strongly on experimental conditions that influence excitability, e.g. extracellular ion levels ([K^+^] and [Ca^2+^] in particular; Somjen (2004)), synaptic activity, temperature, ion channel modulation, pharmacology, and stimulation protocol (Stasheff et al., 1993; Krook-Magnuson et al., 2011; Sheffield et al., 2013; Thuault et al., 2013; Suzuki et al., 2014; Elgueta et al., 2015; Thome et al., 2018). Definitions of ectopic firing have also varied. Here, we included all EAPs in our analyses, and used a modified version of a previous definition of barrage firing in hippocampal NPY cells (Sheffield et al., 2011). Our approach distinguished between scattered firing patterns and more regular barrage (e.g. lines 25 and 46 in Figure 4A), which are likely to have qualitatively different effects on synaptic targets.

The characteristics of EAPs we observed in neocortical PV+ cells resemble those of some non-PV+ cortical interneurons. Sheffield et al. (2011, 2013) and Deemyad et al. (2018) described robust “retroaxonal barrage spiking” in ∼80% of *Htrb5b*-expressing interneurons (NPY-expressing interneurons) from hippocampus and somatosensory cortex. Krook-Magnuson et al. (2011) found that ∼80% of NPY-expressing interneurons (neurogliaform and Ivy cells) in hippocampus could be induced to fire similar EAPs. Suzuki et al. (2014) described ectopic barrage firing (full spikes and spikelets) in >80% of neurogliaform cells from piriform cortex, somatosensory neocortex, and hippocampus.

It is clear that a wide variety of inhibitory and excitatory neurons—perhaps all types of vertebrate neurons—can generate EAPs under certain conditions. Unfortunately, studies of the phenomenon have been sporadic. The conditions that induce ectopic firing, its variable threshold and expression across cells, and its potentially dramatic effects on neuronal and circuit function deserve more attention. Tests for ectopic spiking are rarely, if ever, included in large-scale studies characterizing the intrinsic physiology of neuron subtypes (Markram et al., 2004; Rudy et al., 2011; Gouwens et al., 2020). We suggest that the propensity for generating EAPs should be routinely included in the intrinsic physiological phenotype of neurons.

We have used the term “ectopic” to describe action potentials with the hallmarks of spikes initiated at axonal sites distal to the axon initial segment. EAPs usually arise from membrane potentials more hyperpolarized than the threshold of somatically induced action potentials (Figs. 2,4,6,7). EAP amplitudes can vary categorically (Fig. 5); they occur at wide-ranging latencies following induction (Fig. 3). The precise site(s) of EAP initiation in PV+ cells is unknown. The axons of cortical PV+ cells include many myelinated segments, most prominently near the soma; myelin distribution tends to be patchy, and internodes are relatively short (Peters and Proskauer, 1980; DeFelipe et al., 1986; Micheva et al., 2016; Stedehouder et al., 2017; Stedehouder et al., 2019). EAP initiation sites in PV+ cells could include axon terminals, branch points, unmyelinated segments of axons, and/or nodes of Ranvier. Studies of crustacean neurons using electrophysiology and voltage imaging have pinpointed ectopic spike initiation zones along lengths of unmyelinated axonal trunks (Städele and Stein, 2016; Städele et al., 2018).

The mechanisms of EAPs have been investigated in detail for only a few types of neurons (Pinault, 1995; Bucher and Goaillard, 2011a). EAPs seem to be a pervasive feature of some crustacean neurons (Pinault, 1995; Bucher et al., 2003; Daur et al., 2009; Bucher and Goaillard, 2011; Daur et al., 2019). Neurons of the lobster stomatogastric ganglion are perhaps the best understood. Some lobster motor axons can generate spontaneous EAPs in the absence of central drive, apparently because axonal spike thresholds are close to resting potential and the membrane is regulated by multiple ion channels and transporters—in particular, subtypes of hyperpolarization-activated cyclic nucleotide-gated cation (HCN) channels, voltage- and calcium-activated K^+^ channels, and an electrogenic Na^+^/K^+^ pump. Some HCN channels can be modified by neural activity and neurotransmitters, leading to transient axonal depolarization above threshold. In particular, dopamine can, via an increase in cAMP, enhance axonal HCN conductances and induce tonic EAPs in motor axons (Daur et al., 2009; Bucher and Goaillard, 2011; Ballo et al., 2012; Daur et al., 2019). The generality of this mechanism across neuron types and species is not known, but the available data are suggestive. In the rodent, HCN channels have been shown to enhance AP initiation and increase action potential propagation speed in PV+ basket cells, and are expressed in their axons (Roth and Hu, 2020). Elgueta et al. (2015) showed that barrages of EAPs of PV+ interneurons in the dentate gyrus originate in distal axons, and they implicated axonal HCN channels in EAP generation. HCN channels were also found to be critical in sustaining barrages of EAPs in certain layer 1 (presumably PV-negative) interneurons (Rózsa et al., 2023). Given that HCN channel expression and kinetics change over the course of development (Yang et al., 2018), it is possible that these changes also have a bearing on our finding that EAPs are more readily elicited in older animals.

The ectopic firing of our mouse PV+ cells resembles that of rodent (Sheffield et al., 2011) and human (Chittajallu et al., 2020; Rózsa et al., 2023) NPY cells, so the two interneuron types may share similar mechanisms. EAP barrages can be induced in hippocampal NPY cells by a variety of axonal stimuli, and somatic depolarization is not required. Sheffield et al. (2011) suggested that the axons of NPY cells slowly integrate spikes, respond with EAP barrages, and transmit a signal for barrage spiking to neighboring interneurons. Their mechanistic experiments (Sheffield et al., 2013; Deemyad et al., 2018) imply that ectopic spiking in NPY interneurons requires depolarization and Ca^2+^ accumulation in periaxonal astrocytes. Bidirectional neuro-glial signaling may involve GABA released from NPY axons and astrocytic glutamate acting on metabotropic receptors of NPY axon terminals. These intriguing clues suggest that complex interneuron-glia interactions underlie EAPs.

Our recordings often revealed subthreshold fluctuations of somatic membrane potential, both transient and protracted, that occurred during ectopic spiking (Figs. 6,7). These fluctuations may have originated in axons or terminals, perhaps due to axonal ion channel conductances modulated by the slow integration of stimulus-evoked axonal spikes. The presence of EAPs that occurred both during and outside of subthreshold fluctuations (Figs. 6C, 7A7), and interspersed large and small EAPs (Fig. 8) suggest EAPs originating from two or more distal axonal branches. The small EAPs that we saw, however, could be interpreted differently – as gap junction-mediated spikelets.

Gap junctions electrically couple most inhibitory interneurons in the cerebral cortex, and may play an important role in network synchrony (Gibson et al., 1999; Beierlein et al., 2000; Tamás et al., 2000; Galarreta and Hestrin, 2001b; Galarreta and Hestrin, 2001a; Connors and Long, 2004; Gibson et al., 2005; Long et al., 2005; Hjorth et al., 2009; Vervaeke et al., 2010). We must, therefore, consider the possibility that the low-amplitude spikelet-like events we interpret here as axonally initiated EAPs are, instead, action potentials electronically conducted (via gap junctions) from neighboring interneurons. The evidence suggests that gap junctions are not a major contributor to the events described here. First, gap junction-mediated spikelets recorded in fast-spiking interneurons typically do not exceed about 0.5-1 mV in amplitude (Tamás et al., 2000; Galarreta and Hestrin, 2001a). The small, spontaneous events we report here are distinctly larger, about 2-6 mV. These small presumptive EAPs were also induced by stimulating the soma of the cell from which they were recorded; it is unlikely they were initiated within a neighboring cell. In addition, these small events were often interspersed among full-amplitude EAPs, in-phase with them, and at the same frequency (see Figs. 2E and 4D for examples). It is not clear how gap junctions would mediate such an alternating spike-spikelet pattern. Finally, the shapes of action potentials electrotonically conducted from one PV+ interneuron to another tend to be distinctly biphasic (see experimental and modeling results of Gibson et al., 2005), and do not resemble the shapes of small EAPs. This is a consequence of the singular waveforms of PV+ action potentials (i.e. a rapid, brief spike depolarization followed by a deep, slow afterhyperpolarization) and the inherent low-pass-filtering properties of gap junction-mediated communication.

The functions of EAPs are as mysterious as their mechanisms. Persistent ectopic firing in PV+ cells would generate periods of inhibitory influence independent of further synaptic input. The downstream targets of PV+ cells in cerebral cortex include local pyramidal cells (their somata, proximal dendrites, and axon initial segments) and several types of inhibitory interneurons, including other PV+ cells and SOM cells (Tremblay et al., 2016). The wide variability in ectopic firing patterns across individual PV+ cells (Fig. 3) could have differential inhibitory impacts. In general, persistent EAPs could disrupt the balance of excitation and inhibition in local cortical circuits, and alter the timing, probability, and synchrony of spikes in pyramidal cells and other interneurons. Finally, autonomous EAPs, which functionally disconnect a cell’s axon from its synaptic inputs, could disrupt normal processing related to sensory perception, motor control, and cognition (Pinault, 1995; Keros and Hablitz, 2005; Papatheodoropoulos, 2008; Connors and Ahmed, 2011; Sheffield et al., 2011; Dugladze et al., 2012; Suzuki et al., 2014).

How might ectopic spiking of normal inhibitory interneurons be adaptive? Early studies of seizure-related ectopic spiking in excitatory projection neurons emphasized how this phenomenon could contribute to the scope and propagation of seizures (Gutnick and Prince, 1972; Gutnick and Prince, 1974; Noebels and Prince, 1978; Pinault and Pumain, 1985; Pinault, 1995). However, similar ectopic spiking in interneurons could mitigate the intensity and spread of seizures by shifting the balance of synaptic tone toward enhanced local inhibition. If ectopic spiking occurs in PV+ cells under normal conditions *in vivo*, it is also possible that autonomous axonal spiking might contribute to cortical rhythm generation and synchrony (Sheffield et al., 2011). Cortical PV+ interneurons are extensively coupled by dendritic and somatic electrical synapses (Connors and Long, 2004), which could facilitate the coordination of ectopic spiking across networks of PV+ cells and thereby promote synchrony.

EAPs may have roles in neuropsychiatric disorders beyond epilepsy. They have been detected in multiple pathological states, including epilepsy, neuropathic pain, nerve injury, neuroinflammation, and demyelination (Rosen and Vastola, 1971; Gutnick and Prince, 1972; Pinault, 1995; Hoffmann et al., 2008). Imbalanced excitatory/inhibitory PV+ cell tone is often cited as core features of neurodevelopmental dysfunctions (Yizhar et al., 2011; Vecchia and Pietrobon, 2012; Nelson and Valakh, 2015; Ghatak et al., 2021). Perturbations of ectopic firing in PV+ cells could contribute to a range of neuropsychiatric disorders, and potentially suggest novel therapeutic targets.

## EXPERIMENTAL DESIGN AND STATISTICAL METHODS

### Mice

All procedures were carried out in accordance with the NIH Guidelines for the Care and Use of Laboratory Animals and were approved by the Brown University Institutional Animal Care and Use Committee. Animals were maintained on a 12:12-hr light-dark cycle and provided food and water ad libitum. We examined five strains of animals, which were crosses of the following mice: 1) Parvalbumin (PV)-Cre (The Jackson Laboratory #017320) with Ai14 (a Cre-dependent TdTomato reporter; The Jackson Laboratory #007914); 2) Somatostatin (SOM)-Cre (The Jackson Laboratory #013044) with Ai14; 3) PV-Cre x GCaMP6f (fast Cre-dependent calcium indicator; The Jackson Laboratory #025393); 4) PV-Cre x ChR2-YFP (Cre-dependent channel rhodopsin; The Jackson Laboratory #024109); 5) PV-Cre x Halorhodopsin (Cre dependent inhibitory light-dependent chloride channel; The Jackson Laboratory #014539). All animals were bred by crossing homozygous Cre mice with homozygous reporter mice, resulting in experimental mice that were heterozygous for the indicated genes. Note that mice were coisogenic strains on a C57/BL6 background, and that while the ChR2-YFP and Halorhodopsin mice were not necessary for present study’s goals they were included to maximize use of existing animals and minimize colony size. Mouse ages ranged between P19-61. Both male and female animals were used and were randomly selected for experimentation before sex was determined.

### Slice preparation

Acute parasagittal brain slices containing OFC and S1BF were prepared for *in vitro* recording using methods similar to those described previously (Crandall et al., 2017). Mice were first deeply anesthetized with isoflurane in a closed chamber. When the mouse no longer re-oriented itself after being placed on its back, we removed it from the chamber and decapitated with sharp scissors. Before removing the brain, we reflected the fur over the skull laterally, and briefly transferred the head to cold (∼4° C) oxygenated (95% O_2_, 5% CO_2_) slicing solution to facilitate separation of the skull from the brain surface. We then quickly removed the brain from the skull, taking care not to disrupt the olfactory bulb, which provided a useful landmark for the mediolateral position during slicing. Slicing solution was composed of the following (in mM): 3 KCl, 1.25 NaH_2_PO_4_, 10 MgSO_4_, 0.5 CaCl_2_, 26 mM NaHCO_3_, 10 glucose, and 234 sucrose.

We next blocked the brain bilaterally with a razor blade in the parasagittal plane to provide a sufficient mounting surface, then bisected the block with a midline cut. We then mounted both hemispheres to the stage of a vibrating tissue slicer (Leica VT1200S), and cut them at a thickness of 300 µm, yielding parasagittal slices containing OFC. The parasagittal plane was chosen because electrical stimulation of the white matter just under layer 6 elicited cortical activity that extended nearly to the cortical surface, indicating better intralaminar connectivity than the coronal slice plane (Llano et al., 2009; Theyel et al., 2011). An online version of The Allen Brain Atlas was used to identify the boundaries of the orbitofrontal cortex Immediately after slices were cut, we transferred them to warm (32°C) oxygenated (95% O_2_, 5% CO_2_) artificial cerebrospinal fluid (ACSF) containing, in mM: 126 NaCl, 3 KCl, 1.25 NaH_2_PO_4_, 3 MgSO_4_, 1.2 CaCl_2_, 26 NaHCO_3_, and 10 glucose. After 15-20 min, we moved the holding chamber (containing warmed solution) out of the water bath to room temperature where we continued to incubate the slices for a minimum of 45 minutes before recording.

### *In vitro* recordings

For recordings, individual brain slices were transferred to a submersion chamber and bathed continuously at a rate of 3-5 ml/min with warm (31-33°C), oxygenated (95% O_2_, 5% CO_2_) ACSF containing, in mM: 126 NaCl, 3 KCl, 1.25 NaH_2_PO_4_, 1 MgSO_4_, 1.2 CaCl_2,_ and 10 glucose. For experiments involving blockade of fast glutamatergic receptors we used 100 μM 2-Amino-5-phosphonopentanoic Acid (AP5) and 50 μM DNQX disodium salt (DNQX) (Millipore-Sigma, Burlington, MA). We located areas OFC and S1BF by visualizing slices at 4x magnification using infrared differential interference contrast (IR-DIC) optics, and comparing them against images from the Allen Brain Atlas. Once the region of interest was identified we switched to 40x magnification and patched cells expressing reporters identified under epifluorescence illumination accompanied by a small amount of brightfield illumination. This enabled us to continuously visualize labeled cells while patching them. See Figure 1 for anatomy and an example of a recorded PV+ cell. Whole-cell recordings were done in current-clamp using borosilicate glass micropipettes with a K-based internal solution comprised of, in mM: 130 K-gluconate, 4 KCl, 2 NaCl, 10 HEPES, 0.3 EGTA, 4 adenosine triphosphate (ATP)-Mg, 0.3 guanosine triphosphate (GTP)-Tris, and 14 phosphocreatine-Tris (pH between 7.21 and 7.25, 290-291 mOsm). The initial resistance of the microelectrodes was 5-7 MΩ. Reported recordings were not corrected for a 14.5 mV liquid junction potential (measured to confirm calculation).

Electrophysiological data were acquired with a MultiClamp 700B (Molecular Devices, San Jose, CA) microelectrode amplifier and then digitized at 20 kHz with a Digidata1440A (Axon Instruments) acquisition system and Clampex data acquisition software (pClamp 10, Molecular Devices). All signals were low-pass-filtered at 10 kHz prior to digitizing. During whole-cell recordings, the pipette capacitances were neutralized, and series resistances (typically 15–25 MΩ) were compensated online.

Action potentials were counted using an in-house custom program written to automatically detect spike time, resting membrane potential, threshold, and action potential amplitude (Matlab release 2022b, The Mathworks, Natick, MA). All data were visually inspected to ensure program accuracy, and the program threshold was modified for medium- and small-amplitude EAPs.

Threshold was identified as the maximal value of the second derivative of the trace during the period of time preceding the spike peak. *Action potential amplitude* was measured as the difference from cell’s membrane potential preceding current step injection to the peak of the first evoked spike. In the case of EAPs, the action potential amplitude was calculated as the difference between the peak amplitude and the membrane potential immediately prior to the beginning of the action potential.

Phase plots were calculated and plotted in Excel (Microsoft, Redmond, WA). All statistical analyses were performed in Excel. Figures were generated by copying data from pClamp and pasting them into Adobe Illustrator (Adobe, San Jose, CA). Modifications in Adobe Illustrator were restricted to line thickness enhancements and resizing/cropping data to fit figure panels. All manipulations were performed both on the traces and scale bars concomitantly.

### Ectopic action potential induction

We initially tested a wide array of protocols to determine which was the most effective way to evoke EAPs in the whole-cell current-clamp configuration. These included: 1) evenly spaced one second-long trains of varying frequency; 2) brief (100 µs) current pulses of high amplitude (800-2400 pA), each sufficient to trigger one spike, applied at 60, 100, or 130 Hz to evoke hundreds of spikes over the course of minutes; 2) fixed 1-second long current steps at 800 pA, or 3) a series of increasingly larger amplitudes of one second-long current pulses, starting at intensities below spike threshold for each cell and ending once the cell’s response suggested it was reaching depolarization block. We eventually settled on two suitable protocols: protocols #2 and #3. We found that the both reliably elicited ectopic activity in most cells, and both protocols were used in the present study. Protocol #2 was terminated after persistent ectopic firing was reserved, and protocol #3 was terminated when cells reached depolarization block or fired EAPs constantly until the next current step was applied.

### Coefficient of variation

To quantify the variability of interspike intervals during barrages of ectopic spiking we employed the coefficient of variation (CV_2_) as described in Holt et al. (1996). CV_2_ is obtained by taking the standard deviation of two adjacent ISIs, dividing the result by the mean of the two ISI’s, then multiplying that result by √2. As stated in Holt, et al. (1996): “An average of CV_2_ over *i* estimates the intrinsic variability of a spike train, nearly independent of slow variations in average rate, by comparing adjacent intervals and not widely separated intervals…” CV_2_ is high when ISI variability is high, but in contrast to the traditional coefficient of variation, it is not as influenced by a spike train in which some ISIs are much longer than others.

## Author Contributions

B.B.T. and B.W.C. designed the experiments. B.B.T. and R.J.S. conducted the experiments. B.B.T. and R.J.S. analyzed the results. B.B.T. and B.W.C. wrote the original draft. B.B.T., R.J.S. and B.W.C. edited and finalized the paper.

## Acknowledgements

The authors thank Frederic Pouille, Julia B. Zaltsman, Scott J. Cruikshank, and F. Scott Susi for helpful discussions. We also thank Elizabeth V. McDonnell for preliminary data collection related to this project.

