## Supplemental Table 1 for "Activity-Dependent Ectopic Spiking in Parvalbumin-Expressing Interneurons of the Neocortex"

**Supplemental Information:**

**Supplemental Table 1. Number of action potentials needed to trigger ectopic firing.**

| ***Cell Type*** | ***Action potentials required to elicit first ectopic action potential[1]*** | ***Number of ectopic action potentials evoked in most active run*** |
|---|---|---|
| OFC PV Cells<br>(n = 43) | 1,236±772 triggered action potentials (median 989) | 102.8±149.7 |
| S1BF PV Cells<br>(n = 21) | 1,516 ± 1679 action potentials (median 997) | 64.5 ± 79.6 |
| OFC SOM Cells<br>(n = 7) | 1,808 ± 1310 (median 1004) | 8.4 ± 18.2 |

[1] The number of action potentials evoked during whole-cell patch clamp recording prior to the first EAP: mean +/- SD
